# Shared and distinctive neighborhoods of emerin and LBR revealed by proximity labeling and quantitative proteomics

**DOI:** 10.1101/2022.05.11.491529

**Authors:** Li-Chun Cheng, Xi Zhang, Kanishk Abhinav, Julie A Nguyen, Sabyasachi Baboo, Salvador Martinez-Bartolomé, Tess C Branon, Alice Y Ting, Esther Loose, John R Yates, Larry Gerace

## Abstract

Emerin and LBR are abundant transmembrane proteins of the nuclear envelope (NE) that are concentrated at the inner nuclear membrane (INM). Although both proteins interact with chromatin and nuclear lamins, they have distinctive biochemical and functional properties. Here we have deployed proximity labeling using the engineered biotin ligase TurboID (TbID) and quantitative proteomics to compare the neighborhoods of emerin and LBR in cultured mouse embryonic fibroblasts (MEFs). Our analysis revealed 232 high confidence proximity partners (HCPP) that interact selectively with emerin and/or LBR, 49 of which are shared by both. These included previously characterized NE-concentrated proteins, as well as a host of additional proteins not previously linked to emerin or LBR functions. Many of these are TM proteins of the ER and include two E3 ubiquitin ligases. Using the proximity ligation assay as an orthogonal approach, we validated the interactions described by proximity labeling for 11/12 proteins analyzed, supporting the robustness of our analysis. Overall, this work presents methodology that may be used for large-scale mapping of the landscape of the INM and reveals a group of new proteins with potential functional connections to emerin and LBR.

## Introduction

The nuclear envelope (NE), the membrane system that forms the nuclear boundary, is a sub-domain of ER that compartmentalizes chromosomes and associated metabolism (Dultz and Ellenberg, 2007). It contains inner and outer nuclear membranes joined at nuclear pore complexes (NPCs), the conduits for molecular transport between the nucleus and cytoplasm (Beck and Hurt, 2017; Knockenhauer and Schwartz, 2016; Lin and Hoelz, 2019). The outer nuclear membrane (ONM) is contiguous with the peripheral ER and shares biochemical and functional properties with the latter, whereas the inner nuclear membrane (INM) enriches a distinctive set of proteins (Katta et al., 2014; Pawar and Kutay, 2021). NPCs are ∼100 mDa supramolecular assemblies containing multiple copies of ∼30 different polypeptides (nucleoporins or Nups) that form aqueous channels spanning the NE (Knockenhauer and Schwartz 2016, Beck and Hurt 2017, Lin and Hoelz 2019). NPCs restrict the passive diffusion of molecules larger than ∼20 kDa, and additionally, facilitate the trafficking of nuclear transport receptors and associated cargoes for nucleocytoplasmic movement of most proteins and RNAs.

In higher eukaryotes, the most prominent structural component of the INM is the nuclear lamina (NL), a protein meshwork lining the NE (Burke and Stewart, 2013; Dobrzynska et al., 2016; Gruenbaum and Foisner, 2015; Wong et al., 2021). The backbone of the NL comprises polymers of nuclear lamins, type V intermediate filament proteins (de Leeuw et al., 2018). Most differentiated mammalian cells contain three distinct lamin subtypes: the alternatively spliced lamins A and C, lamin B1 and lamin B2. The INM also contains over 25 widely expressed proteins that are concentrated at the NE (Cheng et al., 2019; Malik et al., 2010; Pawar and Kutay, 2021; Schirmer et al., 2003), most of which are membrane-embedded via transmembrane (TM) segments. Collectively, nuclear lamins and associated proteins have essential roles in the cell nucleus supporting nuclear structure and mechanics (Cho et al., 2017; Maurer and Lammerding, 2019; Miroshnikova and Wickstrom, 2022), chromatin organization and maintenance (Hildebrand and Dekker, 2020; Kim et al., 2019), and regulation of signaling and gene expression (Choi and Worman, 2014; Gerace and Tapia, 2018). Correspondingly, at least 15 human diseases are caused by mutations in NL proteins (Shin and Worman, 2021; Wong and Stewart, 2020).

TM proteins of the INM are synthesized and become membrane-integrated in the peripheral ER. In higher eukaryotes, they are thought to accumulate at the INM largely by a diffusion-retention mechanism, involving passive movement in the plane of the lipid bilayer around NPCs coupled with accumulation at the INM by binding to NL and chromatin or other intranuclear components (Katta et al., 2014; Ungricht and Kutay, 2015). With this mechanism, exchange of TM proteins between ONM and INM is intrinsically bidirectional and is limited by the size of their cytoplasmic/nucleoplasmic domains. The partitioning of TM proteins between the peripheral ER and NE, rather than being an invariant cell feature, can depend on the cell type (Malik et al., 2010) and dynamically change in different functional states (Le et al., 2016). Superimposed on this passive diffusion process, some INM proteins in higher eukaryotes also may deploy receptor and signal-mediated facilitated diffusion around the NPC (Mudumbi et al., 2020) as established in yeast (King et al., 2006; Meinema et al., 2011). Model INM proteins contain multiple regions that promote their accumulation at the NE, presumably due to associations with different cognate binding partners (Berk et al., 2013; Ungricht and Kutay, 2015). Many abundant INM proteins are suggested to occur in heterogeneous and dynamic macromolecular assemblies rather than in discrete complexes of fixed stoichiometry. Biochemical characterization of complexes containing these proteins has been confounded by the resistance of the NL to chemical solubilization. Accordingly, in vivo approaches are needed to further explore the protein interactions of individual INM proteins.

Proximity labeling is a powerful approach to map the local environments of proteins in living cells (Qin et al., 2021; Samavarchi-Tehrani et al., 2020). This method commonly involves ectopic expression of a “bait” protein genetically fused to an engineered biotin ligase (e.g., BioID) or peroxidase (e.g., APEX2), which produces a short-lived reactive intermediate that covalently attaches biotin to “prey” proteins within an ∼10-20 nm radius. Enrichment of biotin-coupled proteins under denaturing conditions followed by mass spectrometry (MS) analysis allows profiling of the protein environment(s) of specific baits. However, prey labeling is affected by many variables, including the level of ectopic bait expression, the duration of biotin labeling and the abundance of the prey themselves (Go et al., 2021; Samavarchi-Tehrani et al., 2020). Moreover, specific prey can have several functions and reside in multiple organelles, making labeling patterns difficult to interpret. Quantitative, comparative analysis of different baits can help assess the significance of prey labeling, although understanding the biological meaning of results requires functional studies.

Here we deployed proximity labeling with TurboID (TbID) probes and quantitative MS to compare the neighborhoods of two abundant TM proteins of the INM, emerin (Emd, UniprotKB P50402) and LBR (UniprotKB Q13749). These proteins, which have been extensively analyzed in mammalian cultured cell models, have been linked to human diseases and implicated in chromatin tethering to the NE (Berk et al., 2013; Olins et al., 2010). Emerin and LBR both contain a nucleoplasmic domain of ∼200 residues harboring folded and intrinsically disordered regions (see Fig. 1). However, they differ in their detailed properties, including their interaction partners and mechanisms for chromatin regulation. The nucleoplasmic domain of emerin (pI ∼ 5.0) contains an ∼40 aa “LEM” (<u>L</u>AP2, <u>e</u>merin, <u>M</u>AN1) homology domain that interacts with the chromatin-associated protein BAF (Berk et al., 2013). By contrast, the amino terminal region of LBR (pI ∼10) interacts with chromatin through at least two separate regions, a chromodomain that binds to heterochromatin proteins HP1-α and HP1-γ (Ye et al., 1997), and a Tudor domain that associates with the H4K20me2 epigenetic mark (Hirano et al., 2012). In addition to chromatin regulation, emerin functions in the peripheral ER as well as at the NE (Le et al., 2016), and LBR plays an essential role in cholesterol biosynthesis through its sterol C14 reductase activity (Tsai et al., 2016).

**Figure 1.**
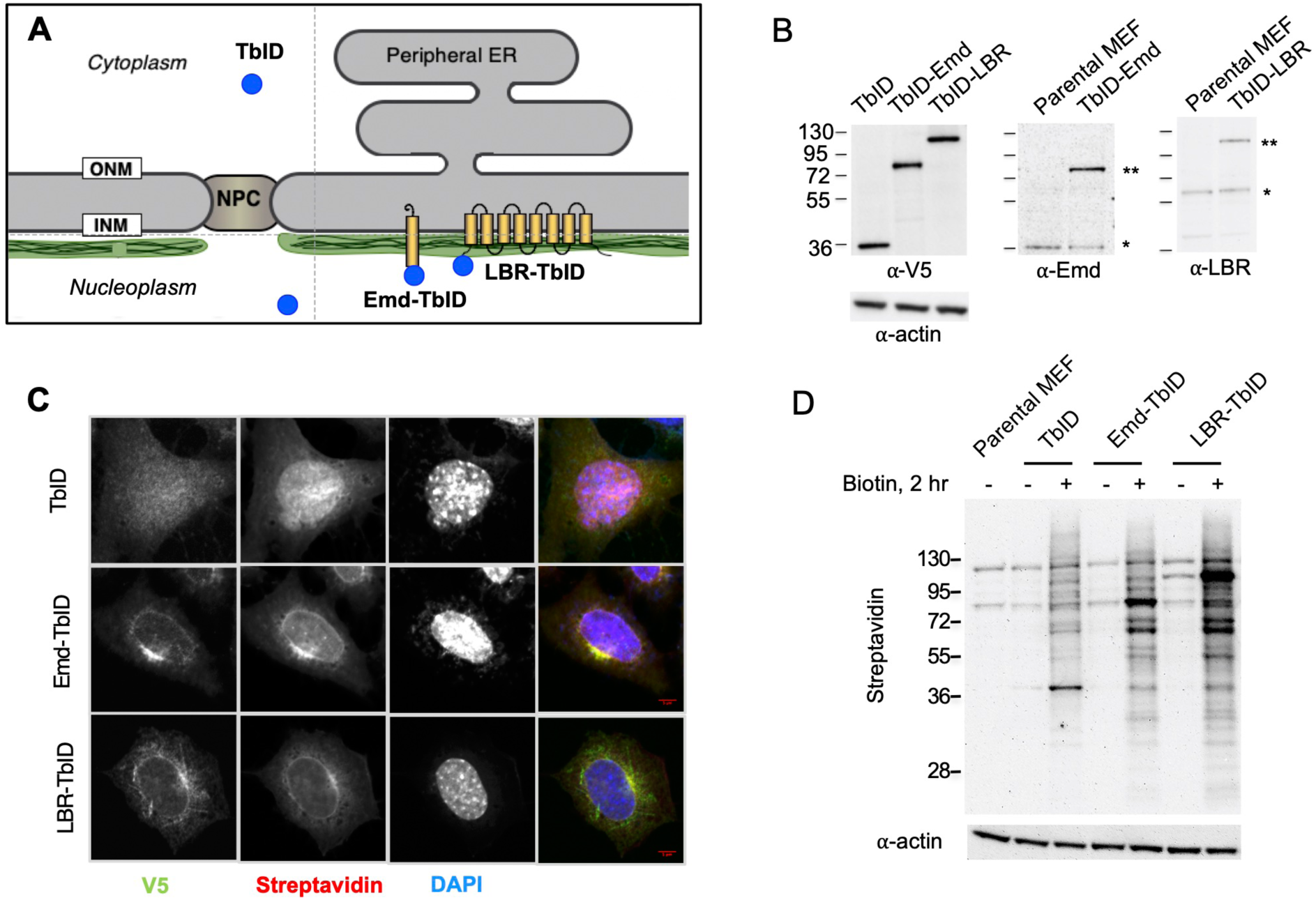
Proximity labeling strategy to investigate the neighborhoods of emerin and LBR using TbID fusions. (A) Schematic diagram of the NE, illustrating the continuity of the INM and ONM at the NPC, and the contiguity of the ONM with the peripheral ER. Ectopically expressed constructs with TbID fused to the N-terminus of emerin (Emd-TbID) or LBR (LBR-TbID) were concentrated at the INM as depicted, but also were located in the peripheral ER and other endomembranes at a lower concentration (not shown). Unfused TbID lacking a TM domain, which served as a control, is distributed throughout the nucleoplasm and cytoplasm. The NL is indicated by green stipple. (B) Western blots of parental MEFs, or MEFs stably expressing V5-tagged TbID, TbID-Emd, or TbID-LBR as indicated. Blots were probed with anti-V5 tag or anti-actin (left panels), anti-emerin (middle panel) or anti-LBR (right panel). (C) Immunofluorescence micrographs of MEFs stably expressing TbID constructs (panel B) that had been incubated with exogenous biotin and stained as indicated to detect the V5 tag, biotin (streptavidin) or DNA (DAPI). Merged images, right panels. Bar, 1 *µ*m. (D) Western blots of parental MEFs, or MEFs stably transduced with TbID constructs as indicated. Cell samples were incubated without (-) or with (+) 500 uM biotin for 2 h prior to probing with streptavidin or anti-actin, as indicated.

Consistent with the biochemical and functional differences between emerin and LBR, our proximity analysis revealed distinctive sets of proteins that were selectively labeled by each bait. In addition, the two baits yielded strong labeling of a shared set of proteins, many of which may reflect a more general INM environment. Using an orthogonal approach, we confirmed proximity relationships for 11 prey sets, including two ubiquitin E3 ligases not previously linked to the NE. Together our results reveal distinctive and shared environments for emerin and LBR and identify new proteins with potential functions at the INM.

## Experimental Procedures

### Cell culture and lentiviral transduction

Mouse embryonic fibroblasts (MEFs), C3H/10T1/2 mouse mesenchymal stem cells (ATCC, CCL-226) and 293T cells (ATCC, CRL-3216) were cultured at 37°C in 5% CO_2_ in DMEM supplemented with 10% FBS, 2 mM L-glutamine, MEM nonessential amino acids, and antibiotics (100 units/ml penicillin and 100 *µ*g/ml streptomycin) (Gibco), termed “standard growth medium”. MEFs were derived in-house from C57BL6/J mice by immortalization with the SV40 T antigen. Cells were passaged at 80-90% confluency and medium was changed every 48 hours. Cultures were routinely checked for mycoplasma contamination.

Lentivirus was produced in 293T cells. Cultures grown to 80% confluency were shifted to standard growth medium without antibiotics, and cells were transfected with a mixture of pRSV-Rev (Addgene # 12253), pMDLg/pRRE (Addgene # 12251), pCMV-VSV-G (Addgene # 8454), and the lentiviral expression plasmid pLV-EF1a (Addgene # 85132) containing the gene of interest, using Lipofectamine 2000 (Thermo Fisher, 11668019). 48 hours post-transfection, the culture medium containing the virus was harvested, cleared by low-speed centrifugation and filtered through a 0.45 *µ*m filter (GE Healthcare Whatman) to yield “lentivirus supernatant”. For lentiviral transduction of MEFs, trypsinized cells were resuspended in standard growth medium without antibiotics and plated in 6-well culture plates (5 × 10^4^ cells/well) after mixing with 10 *µ*g/ml polybrene (EMD Millipore) and lentivirus supernatant. Following 3 days of culture, cells were treated with 3 mg/ml puromycin (Invitrogen) for an additional 3-5 days to select for cells that had integrated the viral DNA. Cell populations were then expanded and frozen.

### Biotin proximity labeling, subcellular fractionation, and streptavidin pulldown

The expression constructs used for proximity labeling were unfused TbID and TbID fused to the N-terminus of emerin (Emd) or LBR. All constructs had an N-terminal V5 epitope tag. The protocol for biotin proximity labeling with TurboID was modified from (Branon et al., 2018). For proteomics analysis, each bait sample comprised two 15 cm plates of stably transduced MEFs at 80-90% confluency. Four independent samples were analyzed for each bait. The standard labeling conditions involved incubation of cells at 37°C for 120 min with 500 *µ*M biotin (Sigma, B4501), diluted from a 100 mM biotin stock solution made in DMSO. Labeling was terminated by transferring culture plates to ice and washing plates 3 times with ice cold PBS. Next, plates were washed 3 times with ice cold homogenization buffer (HB; 10 mM HEPES pH 7.8, 10 mM KCl, 1.5 mM MgCl_2_, 0.1 mM EGTA, 1 mM DTT, 1 mM PMSF, and 1 *µ*g/mL each of pepstatin, leupeptin, and chymostatin). Cells then were swollen by adding 1ml HB buffer to each plate and incubating for 15 min on ice. Subsequently cells were scraped off the plates using a cell lifter (Tradewinds Direct, 70-2180). The scraped cell suspension was disrupted with ∼20 strokes of a tight-fitting Dounce homogenizer, sufficient to release ∼90% of nuclei from the cell bodies. The resulting homogenate was fractionated by layering on top of 0.8 M sucrose cushion in HB and centrifuging at 2000 RPM for 10 min in a Beckman JS-5.2 swinging bucket rotor, yielding a low-speed nuclear pellet and post-nuclear supernatant. The nuclear pellet was resuspended in 1 ml HB and was sonicated with five, 5-sec pulses at 40% vibration amplitude using a Fisher Scientific 60 Sonic Dismembranator. Proteins in the nuclear pellet were solubilized by adding SDS to 2% and incubating at 95°C for 5 min. Insoluble aggregates were removed by centrifugation at 20000g for 20 min, and the supernatant was diluted to 0.2% SDS with water. Biotinylated proteins were enriched with 50 *µ*L per sample of streptavidin conjugated Dynabeads (MyOne Streptavidin C1, 65001), by incubating for 2 hours on a rotating wheel at room temperature. After pulldown, beads were washed 5 times with 8 M urea. After the final wash beads were resuspended in 8 M urea and were subsequently processed for proteomics analysis as below.

### Preparation of peptide digest for proteomics

The streptavidin Dynabeads (above) were resuspended and washed twice in 10 mM EPPS (N-(2-Hydroxyethyl)piperazine-N′-(3-propanesulfonic acid)) pH 8.5. Next, 5 or 10% of the beads were removed for quality control using SDS PAGE and Western blotting, and the remaining 90 or 95% were used for digestion. Buffer was exchanged into 8 M urea in 10 mM EPPS pH 8.5, 20 *µ*L. The sample was reduced with 10 mM TCEP at room temperature (RT) for 30 min, alkylated with 10 mM iodoacetamide at RT for 30 min in the dark, and then diluted 8-fold to 1 M urea using 10 mM EPPS pH 8.5. To digest, 2 *µ*g Lys-C/Trypsin mix protease (Promega, Mass spec grade) were added to each sample (10 mM EPPS pH 8.5, 1 mM CaCl_2_). The mixture was shaken at 800 rpm at 37 °C overnight, centrifuged and magnetically separated to recover the digested supernatant. As a quality control for protein digestion efficiency, 5 or 10% of the supernatant was acidified with 3% (m/v) formic acid and analyzed by LC-MS/MS. The digest was then stored at -80 °C or immediately labeled with TMT. All concentrations are final values unless noted otherwise.

### TMT labeling and peptide fractionation

To quantitatively compare the four replicates of samples from the TbID, TbID-Emd and TbID-LBR constructs, plus samples from an additional two TbID constructs not considered in this study, TMT 11-plex isobaric labels (Thermo Fisher, A34808, A34807) were used to prepare two sample sets for each LC-MS run. Each 11-channel set comprised two replicates of the five constructs and one common reference channel. The reference channel contained the equal-portion mixture from all sample replicates and was used to normalize peptide quantity between the two runs.

The peptide BCA assay was performed on streptavidin enriched, protease-digested samples (above section) following the manufacturer’s manual (Thermo Fisher, 23275). The initial analysis showed that peptide amounts were low, so an equal portion of the total sample was used for each TMT labeling reaction in subsequent experiments. Each 0.8 mg vial of TMT reagent was dissolved with 44 *µ*L anhydrous acetonitrile, yielding four aliquots of 0.2 mg (11 *µ*L each), and was used within 5 min or temporarily stored at -80°C. Each peptide solution was mixed with 30% (v/v) acetonitrile and reacted with 0.2 mg TMT label solution at RT for 60-80 min. To check labeling efficiency, 2 or 5 μL of each channel was retrieved, quenched with 0.3% NH_2_OH, pooled at equal volumes, and analyzed by LC-MS/MS for %TMT labeling, while the remainder of the sample was stored at -80 °C. Once labeling efficiency exceeded 95%, TMT samples were quenched with 0.3% (m/v) NH_2_OH at RT for 15-20 min, acidified with 3% formic acid to ∼pH 2.5, pooled, and vacuum centrifuged to remove acetonitrile. The samples were then desalted with a C18 peptide desalting spin column (Thermo Fisher, 89852).

To deepen LC-MS data acquisition, TMT-labeled peptides were pre-fractionated with the basic pH reversed-phase C18 peptide fractionation kit following manufacturer’s manual (Thermo Fisher, 84868). Typically, TMT peptides were redissolved in buffer A (0.1% formic acid, 5% acetonitrile in H_2_O), loaded to pre-conditioned high pH fractionation spin column, washed with H_2_O, then with high pH 5% acetonitrile to remove excessive TMT labels, and eluted at high pH with increasing gradient of acetonitrile into 7-10 fractions. Each fraction was vacuum centrifuged to remove acetonitrile, and redissolved in 20 *µ*L buffer A. An autosampler was used to inject 10 *µ*L of each fraction into LC-MS.

### Dimethylation labeling

The 3-plex dimethylation quantitation was used to analyze the nuclear pellet fraction of Emd-TbID MEFs to compare protein capture on streptavidin beads as a function of the biotin concentration and labeling time (50 μM vs 500 μM biotin; 10 min, 1 hour, 2 hours). Isotopic formaldehyde/NaBH_3_CN methylates the free amine groups at the N-terminus and Lys side chains, and quantitation is based on the relative MS peak intensity of the isotopic versions of the common peptides. The 3-plex dimethylation contained three isotopic channels: light (L, COH_2_, NaBH_3_CN, +28.0313 Da), medium (M, COD_2_, NaBH_3_CN, +32.0564 Da) and heavy (H, ^13^COD_2_, NaBD_3_CN, +36.0757 Da). To compare 6 labeling conditions and constructs, 4 sets of 3-plex mixtures were prepared for LC-MS. The L and M were used for to compare two conditions. The H channel was used as the reference and contained an equal-portion mixture from all original samples. Typically, 50 μL of peptide solution in 10 mM EPPS pH 8.0 were mixed with 4 μL of freshly made 4% (m/v) CH_2_O or CD_2_O and 4 μL of 0.6 M NaBH_3_CN or NaBD_3_CN, and incubated at RT for 1 hour. The samples were then quenched with 15 μL of 0.2 M NH_4_HCO_3_, acidified with final 5% (m/v) formic acid, pooled as 3-plex mixtures and vacuum centrifuged to remove acetonitrile. The peptide samples were then desalted with C18 desalting tips (Thermo Fisher, 84850) and injected into LC-MS.

### Mass spectrometry data acquisition

TMT-labeled peptides were analyzed using an EASY-nLC 1200 UPLC coupled with an Orbitrap Fusion mass spectrometer (Thermo). LC buffer A (0.1% formic acid, 5% acetonitrile in H_2_O) and buffer B (0.1% formic acid, 80% acetonitrile in H_2_O) was used for all analyses. Peptides were loaded on a C18 column packed with Waters BEH 1.7 μm beads (100 μm × 25 cm, tip diameter 5 μm), and separated across 180 min: 1-40% B over 140 min, 40-90% B over 30 min and 90% B for 10 min, using a flow rate of 400 nL/min. Eluted peptides were directly sprayed into MS *via* nESI at ionization voltage 2.8 kV and source temperature 275 °C. Peptide spectra were acquired using the data-dependent acquisition (DDA) synchronous precursor selection (SPS)-MS3 method. Briefly, MS scans were done in the Orbitrap (120k resolution, automatic gain control AGC target 4e5, max injection time 50 ms, m/z 400-1500), the most intense precursor ions at charge state 2-7 were then isolated by the quadrupole and CID MS/MS spectra were acquired in the ion trap in Turbo scan mode (isolation width 1.6 Th, CID collision energy 35%, activation Q 0.25, AGC target 1e4, maximum injection time 100 ms, dynamic exclusion duration 10 s), and finally 10 notches of MS/MS ions were simultaneously isolated by the orbitrap for SPS HCD MS3 fragmentation and measured in the Orbitrap (60k resolution, isolation width 2 Th, HCD collision energy 65%, m/z 120-500, maximum injection time 120 ms, AGC target 1e5, activation Q 0.25).

Dimethyl-labeled peptides were analyzed using an EASY nLC 1200 UPLC coupled with a Q Exactive Orbitrap mass spectrometer (Themo). Peptides were directly loaded on to a C18 capillary column packed with Waters BEH 1.7 μm C18 beads (100 μm × 25 cm, 5 μm tip), and separated across 240 min: 1-35% B over 180 min, 35-80% B over 40 min, 80% B for 5 min, then 80-1% B over 5 min and equilibrated with 1% B for 10 min, using a flow rate of 300 nL/min. MS spray voltage was 2.5 kV and capillary temperature 250 °C. Mass spectra were acquired using a DDA10 HCD MS/MS method, where an MS scan (70k resolution, AGC target 1e6, maximum 60 ms, m/z 400-1800) was followed by HCD MS/MS scans of the top 10 most intense precursors with charge states of 2 or higher (isolation window 2 Th,15k resolution, NCE 25, AGC target 1e5, maximum 120 ms, dynamic exclusion of 15 s).

### Mass spectrometry data analysis

Spectra were analyzed using the Integrated Proteomics Pipeline (IP2) platform (IP2, Bruker Scientific LLC). MS/MS spectra were searched using the ProLuCID algorithm (Xu et al., 2015) against a UniProt SwissProt *Mus musculus* reviewed proteome sequence database appended with the sequences of common contaminant proteins, and with the reverse sequences as decoy (UniProt, accessed 2018-11-28, total 34,189 protein entries). Peptide MS/MS spectra (CID for TMT-labeled and HCD for dimethyl-labeled) were searched and filtered using the following parameters: static modifications for TMT (+229.1629 Da; N-term and Lys), dimethyl-tags (light +28.0313 Da, medium +32.0564 Da, heavy +36.0757 Da; N-term and Lys) and carbamidomethylation (57.02146 Cys); dynamic oxidation (+15.9949 Met); dynamic phosphorylation for dimethyl-tag experiment (+79.96633 Ser/Thr/Tyr); precursor mass tolerance 30 ppm; fragment ion mass tolerance, 600 ppm for TMT and 50 ppm for dimethyl-tag; at least 1 tryptic end (Lys/Arg); up to 3 missed cleavages; and minimum peptide length = 4 amino acids. Protein identification required at least 1 peptide (2 peptides for dimethyl-tag) identified per protein, and was filtered to protein false discovery rate < 1% using a target-decoy algorithm (Peng et al., 2003) performed by DTASelect2 (Tabb et al., 2002) in IP2. Quantitation in the TMT experiments was based on reporter ion intensity in MS3 and was performed using Census2 (Park et al., 2008). Dimethyl-labeled peptide quantitation was analyzed by Census2 (Park et al., 2008) by 1) precursor peak ratio of light-, medium-and heavy-versions of the peptide, and 2) spectral counts (NSAF) extracted from individual searches for static Lys-dimethylation of light, medium and heavy versions.

### Statistical and bioinformatic analysis

MS3 intensity values for proteins enriched by the three TbID baits was normalized between the six separate experimental replicates using the V5 tag peptide. After normalization, the relative intensity values of individual entries in a specific TMT channel was expressed as a fraction of the summated intensities in the channel for all detected entries having at least one unique peptide. Normalized intensities from the six replicates were compared by Pearson’s correlation analysis, with a cutoff of 0.1 used for dataset selection. This resulted in elimination of two (unfused) TbID datasets (Supporting Information Table S2). The Student’s 2-tailed T test was used to determine the statistical significance of differences between the normalized intensity values for proteins enriched with Emd-TbID and LBR-TbID (6 replicates) as compared to TbID (4 replicates). GOslim analysis of Biological Process and Molecular Function was performed using Webgestalt (Liao et al., 2019) to determine overrepresentation of terms in the *M. muscularis* database. Analysis parameters involved a FDR < 0.05, use of Affinity Propagation for redundancy reduction, and minimum and maximum identifications set to 5 and 2000, respectively.

### Molecular cloning

Molecular cloning of cDNAs utilized a library generated from C3H/10T1/2 cells. RNA was extracted using TRIzol (Thermo Fisher, 15596026) according to the manufacturer’s instructions. cDNA was synthesized using the iScript cDNA Synthesis Kit (Bio-Rad, 1708890). The ORF of the gene of interest was amplified by PCR using Q5 High-Fidelity DNA Polymerase (New England Biolabs, M0493). TurboID was generated in the Ting laboratory (Branon et al., 2018). DNA fragments, after isolation by agarose gel electrophoresis, were assembled in the pLV-EF1a-IRES-Puro lentivirus backbone (Addgene # 85132), or in lentivirus backbones derived from the latter (pLV-Ef1a-V5-LIC-IRES-Puro; Addgene #120247 or pLV-Ef1a-LIC-V5-IRES-Puro; Addgene #120248). Vector DNA was linearized by digestion with restriction enzymes (New England Biolabs), and constructs were assembled using the NEBuilder HiFi DNA Assembly Kit (New England Biolabs, E2621), in a reaction conducted at 50°C for 20 min. NEB stable *E. coli* cells (New England Biolabs, C3040H) were transformed using 1 μl of NEBuilder reaction mix by heat shock at 42°C for 30 sec. Transformed *E. coli* were selected using ampicillin antibiotic selection and were grown in Luria-Bertani media. Plasmid DNA from individual colonies was extracted using the Monarch Plasmid Miniprep Kit (New England Biolabs, T1010L). All cDNA clones were verified by complete DNA sequencing of the ORF in both the 5’-3’ and 3’-5’ directions.

### RNAi

Depletion of Cgrrf1 and Rnf185 for functional analysis was accomplished with SMARTpool ON-TARGETplus siRNAs (Horizon Discovery: Cgrrf1, L-047570-01-0005; Rnf185, L-064072-01-0005; non-targeting (control), D-001810-10-05). A stock solution was prepared by dissolving the siRNAs in 1X siRNA buffer (300mM KCl, 30mM HEPES pH 7.5, 1.0mM MgCl_2_) to a final concentration of 10 *µ*M. This was further diluted by adding 50 *µ*L of siRNA stock to 950 *µ*L of serum-free standard growth medium to a final concentration of 50nM. A separate mixture was prepared by adding 25 *µ*L DharmaFECT-1 reagent DF1 (Horizon Discovery, T-2001-03) to 1 mL serum-free standard growth medium. A master mix was prepared by adding the DF1 mixture to the siRNA mixture, mixing well, and incubating in the dark at room temperature for 20 min. Meanwhile, MEFs were trypsinized and plated at a density of 0.5 × 10^6^ cells per 10 cm dish. The preincubated master mix was added dropwise to cells and swirled for even distribution. Cells were cultured for 48 hr, at which point the medium was replaced with fresh growth medium. The 48 hr timepoint cells were harvested by trypsinization, and the cell pellet was frozen for subsequent qPCR and Western blot analysis. At 96 hr post-transfection, the second batch of cells was harvested and frozen.

### qRT-PCR

Harvested samples were lysed with 1mL TRIzol (Invitrogen, 15596026) per 2.0 × 10^6^ cells. The lysate was processed following the TRIzol manual to yield an RNA/ethanol mixture. The RNA/ethanol mixture was transferred to a miniprep kit RNA purification column (NEB, T2010S), and RNA was isolated according to the manufacturer. cDNA synthesis was performed by adding 2uL 5X iScript reaction mix (Bio-Rad, 1708889) to 50ng RNA, to a total volume of 10uL. The reaction mix was incubated in a PCR machine using the settings found in the iScript protocol. The primer pairs used for qRT-PCR analysis were 5’-AACCCAGTTCAGCACAAGAGC-3’ and 5’-TCAAGGCCATGCCTGTTGCTA-3’ for Cgrrf1; and 5’-CAGCACCTTTGAGTGCAACA-3’ and 5’-ACTGATGTAAACACGGCCAAC-3’ for Rnf185. SYBR green PCR Master Mix (Applied Biosystems, 4309155) was used, following manufacturer’s instructions. Data was analyzed by calculating dC_t_ values, in which the C_t_ of the housekeeping gene was subtracted from the C_t_ of the gene of interest. Then ddC_t_ values were calculated by subtracting the C_t_ of the control from the dCt. Finally fold change was calculated using 2^(-ddCt).

### Western Blotting

Protein extracts for Western blotting were prepared by sonicating cell pellets on ice in PBS supplemented with protease inhibitors (Thermo Fisher, A32955), 10 mM DTT and 10 mM EDTA, using 40% vibration amplitude with 5, 5-sec pulses. Proteins were then denatured by boiling in sample buffer (2% SDS, 50 mM Tris 6.8 pH, 10% w/v Glycerol, 0.01% w/v bromophenol blue, 2 mM EDTA) at 95°C for 10 min. Protein electrophoresis was done using 4-12% Novex Tris-Glycine gel (Life Technologies) in FAST Run Buffer (Thermo Fisher, BP881). Proteins were then transferred to a nitrocellulose membrane (Thermo Fisher, 10600015) at 24V for 3 hours in transfer buffer at 4°C. Protein transfer was assessed by staining of membranes in Ponceau S solution (0.1% Ponceau S in 5% acetic acid) for 2 min and subsequently de-staining by washing with TBS-Tw20 (Tris-buffered saline with 0.1% Tween-20). Membranes were then blocked with 5% bovine serum albumin in TBS-Tw20 for 1 hour at room temperature and washed three times with TBS-Tw20. Next, blots were labeled by incubating overnight at 4°C with the primary antibody diluted in 0.5% BSA in TBS-Tw20. After washing membranes three times in TBS-Tw20, blots were incubated for 1 hour at room temperature with an HRP-coupled secondary antibody diluted in TBS-Tw20. Finally, membranes were washed three times with TBS-Tw20 and incubated with chemiluminescent substrates solution (Thermo Fisher, 1863059) for 4 min. Chemiluminescence was captured using UVP Biospectrum 810 imaging system.

The primary antibodies and dilutions used for Western blotting were: mouse anti-V5 (Thermo Fisher, R960-25), 1:5000; rabbit anti-emerin (Leica Microsystems, NCL-emerin), 1:2000; and anti-LBR (in-house produced guinea pig antiserum to recombinant human LBR, aa1-218), 1:1000. The secondary antibodies and dilutions were: HRP conjugated goat anti-mouse IgG (Jackson ImmunoResearch, 115-035-003) 1:10000; HRP conjugated donkey anti-rabbit IgG (GE Healthcare, NA934V), 1:10000; and HRP conjugated goat anti-guinea pig IgG (Thermo Fisher) 1:20000. For detection of biotin-labeled proteins (Fig. 1), blots were incubated with HRP conjugated streptavidin (GE Healthcare # RPN1231V), 1:5000.

### Fluorescence microscopy

For cell imaging by immunofluorescence microscopy, cells were plated on sterile cover slips in a 24 well plate in standard growth medium, and were grown for 24 hours. Cells were fixed in 2% paraformaldehyde (Electron Microscopy Sciences, 15710) in PBS for 20 min at room temperature and washed three times with PBS. Blocking and permeabilization involved incubating coverslips with PBS containing 5% goat serum and 0.5% Triton X-100 for 15 min at room temperature. Cells then were washed with PBS containing 0.1% Triton X-100 (PBS-Tx100) three times for 5 min each. Next coverslips were incubated overnight at 4°C with primary antibodies diluted in PBS containing 1% goat serum and 0.1% Triton X-100. The primary antibodies and dilutions were: mouse anti-V5 (Thermo Fisher, R960-25), 1:2000; and affinity purified rabbit anti-lamin B1 (made in-house to residues 391-428 of human lamin B1, Ref. x), 1 *µ*g/ml. After incubation with primary antibodies, coverslips were washed with PBS-Tx100 six times for 5 min each and were incubated with secondary antibodies diluted in PBS-Tx100 for 1 hour at room temperature in the dark. Secondary antibody and dilutions were: goat anti-mouse IgG Alexa Fluor 488 (Thermo Fisher, A28175), (1:1000); and goat anti-rabbit IgG Alexa Fluor 647 (Thermo Fisher, A32733), 1:1000. Coverslips were washed with PBS-Tx100 six times for 5 min each, counterstained with DAPI (1:5000), mounted on slides with ProLong Glass Antifade Mountant (Thermo Fisher, P36980), and sealed using nail polish. In cases were biotin labeling by bait proteins was examined by fluorescence microscopy, cells that had been incubated for 120 min with 500 *µ*M biotin were fixed and permeabilized as described above and were incubated with Alexa Fluor 647 streptavidin (Thermo Fisher, S21374) diluted 1:750, prior to being mounted. Fluorescence imaging was done using Zeiss LSM780 confocal microscope system running Zen software.

### Proximity Ligation Assay

The PLA assay utilized the Duolink system (Sigma-Aldrich). MEFs stably transduced with Myc tagged bait constructs (Myc-MBP, Myc-emerin or Myc-LBR) were seeded at 5,000 cells per well in the 24-well plate on the gelatin coated coverslips. After overnight growth, they were incubated with lentivirus carrying V5-tagged prey expression constructs for 48 hours with the presence of 10 *µ*g/mL polybrene in growth medium. Transduced cells were fixed as described above and then blocked and permeabilized for 15 minutes with PBS containing 5% donkey serum and 0.5% Triton-X. Fixed cells were then incubated with mouse anti-V5 (1:1000, Thermo Fisher, R96125) and rabbit anti-Myc (1:1000, Abcam, ab9106) at 4°C overnight. After antibody incubation, samples were washed three times with PBS and twice with buffer A (10mM Tris, pH 7.4, 150mM NaCl, 0.05% Tween) before annealing with PLA MINUS and PLUS probes (1:10 dilution in antibody diluent provided by Sigma-Aldrich) for 90 minutes at 37°C. After three washes with buffer A, PLA probes were then ligated with Duolink ligase (1:40 in ligation buffer provided by Sigma-Aldrich) for 30 minutes at 37°C, washed again with buffer A, and signals were amplified by polymerization of Texas Red conjugated dNTPs mixture in Duolink amplification buffer (provided by Sigma-Aldrich). After PLA signals were developed, samples were washed twice with buffer B (200mM Tris, pH 7.5, 100mM NaCl) for 10 minutes and once with diluted buffer B (2mM Tris, pH 7.5, 1mM NaCl) for 1 minute. Samples were then equilibrated in PBS and incubated 1 hour with anti-mouse Alexa flour 488 (1:2000, Thermo Fisher) and anti-rabbit Alexa flour 647 (1:2000, Thermo Fisher), washed three times with PBS and counter stained with DAPI before mounted as previously described. Confocal images were done using Zeiss LSM 880 system running Zen software. Typically 10-30 cells were analyzed for each experimental condition.

## Results

### Emerin and LBR neighborhoods in MEFs revealed by proximity labeling

We deployed TbID (Branon et al., 2018), a highly active derivative of BioID (Roux et al., 2012), to probe the neighborhoods of emerin and LBR in cultured cells. Biotin labeling with TbID can be accomplished with a substantially shorter incubation than the 18-24 hours commonly used to analyze BioID samples (Branon et al., 2018; May et al., 2020). Correspondingly, a short labeling protocol with TbID could potentially favor the detection of higher affinity prey. We prepared mouse embryonic fibroblasts (MEFs) that were stably transduced with unfused TbID, or with fusion proteins containing TbID attached to the N-terminus of emerin or LBR (designated Emd-TbID and LBR-TbID, respectively). The TbID constructs migrated at the expected sizes on SDS gels, with expression at ∼1-3 times the levels of the endogenous proteins (Fig. 1B). Whereas endogenous emerin and LBR are concentrated at the INM at steady state (Fig. 1A), they also have peripheral ER pools (Berk et al., 2013; Giannios et al., 2017). Similarly, the emerin and LBR TbID fusions were concentrated at the NE, and also were localized at variable levels to cytoplasmic regions occupied by peripheral ER and to a juxtanuclear region reminiscent of Golgi (Fig. 1C; Supporting Information Fig. S1). By contrast, the unfused TbID was localized diffusely throughout the cytoplasm and nuclear interior (Fig. 1C and Supporting Information Fig. S1). The overall distribution of biotinylated prey was similar to that of the baits, as expected (Fig 1C).

Streptavidin blots of cells expressing the TbID probes revealed strongly increased biotin labeling at 2 hours as compared to the background without exogenous biotin (Fig. 1D), validating our probes and selective labeling with exogenous biotin. We used semi-quantitative proteomics to compare the level of streptavidin enrichment of abundant NE and peripheral ER proteins in Emd-TbID cells over a 10-min to 2-hour biotin labeling time course (Supporting Information Fig. S2 and Supporting Information Table S1). Most NE markers showed progressively increased enrichment over the 2-hour period (up to ∼5-fold). By contrast, most peripheral ER markers showed little or no increase in enrichment after 10 min, suggesting that labeling of these targets by the relevant bait pool was saturated during the 10-min period. These results suggested that a selective increase in labeling of predicted proximity partners (i.e., NE proteins) could be obtained with 2 hour labeling. Therefore, we decided to implement this condition for analysis of the three baits.

The overall workflow for our analysis is depicted in Fig. 2A. After biotin labeling of the three MEF strains, cells were homogenized, and a low-speed pellet containing nuclei and associated/trapped cytoplasmic membranes was prepared. The pellet was solubilized in SDS, and biotinylated proteins were enriched on streptavidin beads. After peptide digestion and labeling for TMT-11, samples were analyzed by quantitative MS. We carried out four independent cell labeling experiments, and analyzed two of these with two technical repeats. This yielded six separate datasets that collectively identified over 2500 proteins (Supporting Information Table S2). To narrow our focus to “high confidence proximity prey” (HCPP), we filtered the datasets to include only proteins that were detected by at least two unique peptides in four or more of the datasets (Fig. 2A). In a second filtering step, we selected proteins that showed at least 3-fold increased enrichment by one or both of the NE baits as compared to unfused TbID (p < 0.05) (Supporting Information Table S2). Some proteins that interact with NE components but are distributed throughout the nuclear interior, such as BAF (Sears and Roux, 2020), could be eliminated by this filtering step.

**Figure 2.**
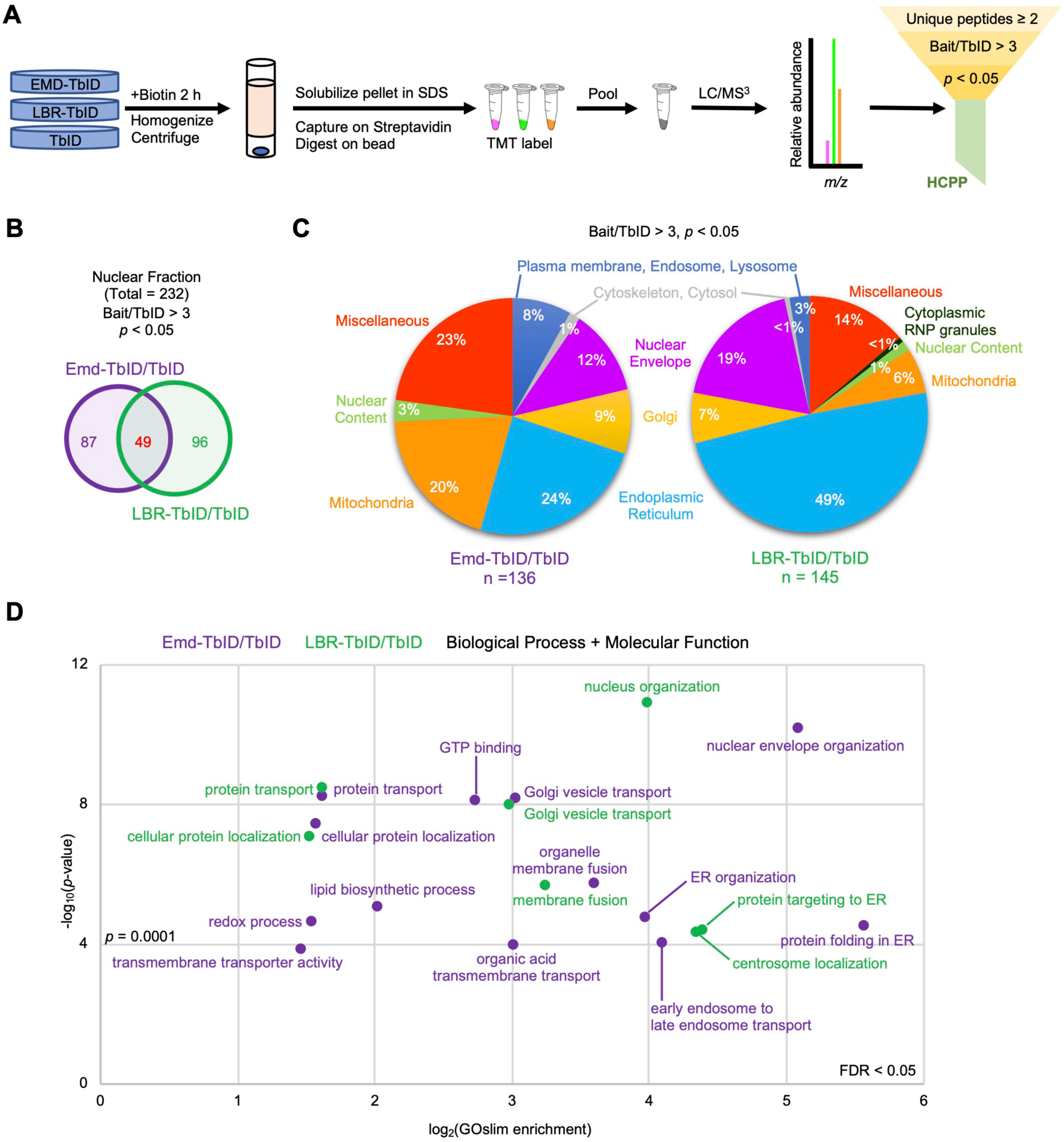
Summary of results from analysis of proximity samples by TMT labeling and proteomics. (A) Workflow depicting steps in the analysis (see text for details). HCPP are streptavidin-enriched proteins from the bait-TbID samples that were detected with at least two unique peptides and that showed at least 3-fold enrichment with p < 0.05, in comparison to unfused TbID samples. (B) Venn diagrams illustrating HCPP selectively labeled by Emd-TbID or LBR-TbID, and overlap between the groups. (C) Pie charts depicting the subcellular locations of HCPP labeled by Emd-TbID or LBR-TbID. (D) Gene annotation summaries (Biological Process and Molecular Function) of HCPP labeled by Emd-TbID or LBR-TbID.

Overall, the analysis detected 232 HCPP. Among these, 136 and 145 prey were enriched with Emd-TbID and LBR-TbID, respectively, including 49 proteins enriched with both probes (Fig. 2B). The great majority of these are TM proteins (Supporting Information, Table S2), and are localized to either the NE, the ER, downstream membranes in the secretory pathway, or membrane organelles known to contact the ER (e.g., mitochondria and the plasma membrane) (Fig. 2C). Correspondingly, the HCPP list for emerin and LBR was strongly enriched for GO functional annotations associated with these organelles, including organization of the NE, nucleus and ER, protein targeting/folding in the ER, vesicular trafficking through the secretory pathway, and lipid biosynthesis (Fig. 2D). As expected, the HCPP group contained well-established “benchmarks” concentrated at the NPC or INM/NL (Supporting Information Table S2, Fig. 3). In addition, it included proteins not evidently concentrated at the NE but nonetheless implicated in NE functions. Examples are the deacetylase Sirt2 (Kaufmann et al., 2016), Ankle2 (Asencio et al., 2012), Reep3/4 (Schlaitz et al., 2013), Lunapark (Lnpk) (Casey et al., 2015; Hirano et al., 2020) and Dnajb12 (Goodwin et al., 2014). However, most of the HCPP have not been previously connected to discrete NE functions.

**Figure 3.**
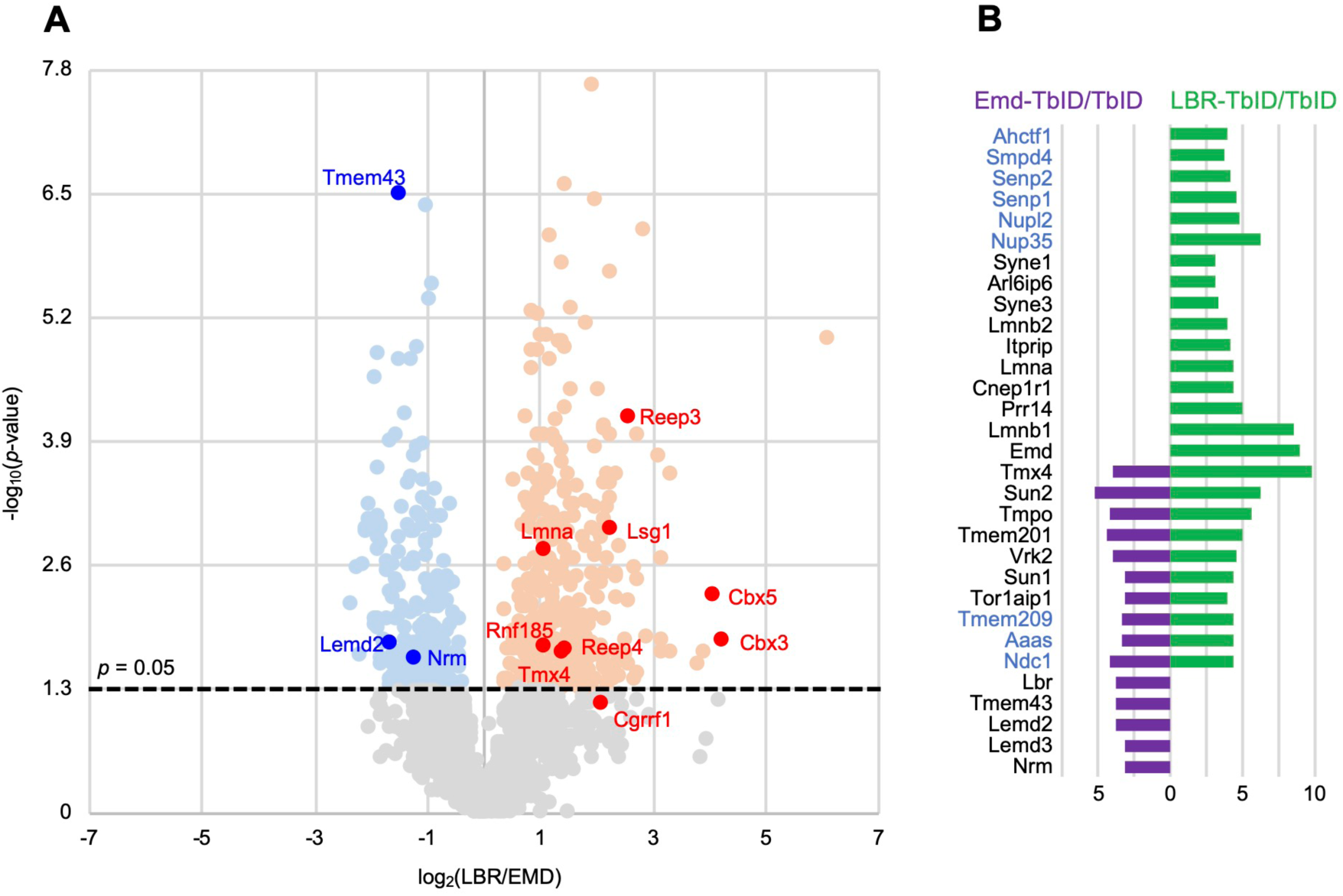
HCPP proteins labeled by Emd-TbID or LBR-TbID. (A) Volcano plot describing preferential labeling of HCPP by Emd-TbID vs LBR-TbID. Prey analyzed in Figs. 4 and 5 are labeled. (B) HCPP from the group of well-characterized NE proteins. Fold-enrichment of hits from Emd-TbID (purple) or LBR-TbID (green) samples, relative to unfused TbID, is indicated on the bottom. Left: NPC proteins are depicted in blue; NL proteins in black.

A volcano plot illustrates that more HCPP were enriched preferentially by the LBR-TbID bait than by Emd-TbID (Fig. 3A). Consistently, 85 HCPP were enriched by > 5-fold by LBR-TbID, whereas only 23 were enriched to this level by Emd-TbID (Supporting Information Table S2). NE constituents in the HCPP group (Fig. 3B) included 14 proteins enriched with Emd-TbID, 25 with LBR-TbID, and 10 with both baits. The NE proteins preferentially enriched with Emd-TbID included the direct emerin-binding proteins Tmem43 (Bengtsson and Otto, 2008) (3.8-fold higher enrichment) and Lemd3/MAN1(Mansharamani and Wilson, 2005) (3.2-fold higher enrichment). Conversely, proteins that were preferentially enriched with LBR-TbID included the LBR interactor lamin B1 (4.8-fold higher enrichment) and two major heterochromatin proteins known to directly associate with LBR, HP1-α (Cbx5, 17-fold higher enrichment) and HP1-γ (Cbx3,19-fold higher enrichment) (Ye et al., 1997). Unsurprisingly, certain NE proteins were enriched strongly by both baits (e.g., Sun1 and Sun2).

### Validation and further analysis of proximity partners

We used the proximity ligation assay (PLA) (Weibrecht et al., 2010) as an orthogonal approach to validate the associations of representative HCPP with relevant bait(s) at the NE (Figs. 4 and 5). This method provides a snapshot of bait-prey associations in cells at steady state. By comparison, another common method to interrogate protein proximity, bimolecular fluorescence complementation (Miller et al., 2015), can overrepresent the steady state abundance of transient interactions because interacting partners become irreversibly trapped in a stable complex. To implement the PLA (Supporting Information Fig. S4), we used populations of MEFs stably transduced with Myc-epitope-tagged versions of either emerin, LBR, or the control maltose binding protein (MBP). Cells were transiently transduced with V5-tagged prey, and after fixation, PLA signal was quantified at the NE/nucleus (Experimental Procedures, Supporting Information Figs. S4A and S4C). We did not analyze the PLA signal in cytoplasmic regions that was seen for some prey, as this may not accurately reflect proximity relationships at the NE. PLA signal was depicted separately for cells with “low” and “high” levels of prey expression, representing the lower and upper half of V5 labeling intensities (Supporting Information Fig. S4A). Using the well-established Cbx5/LBR interaction as a calibration model, we found a roughly linear correlation between PLA signal and level of bait and prey expression over the expression range analyzed (Supporting Information Fig. S4D), supporting the validity of both low and high prey expression datasets.

**Figure 4.**
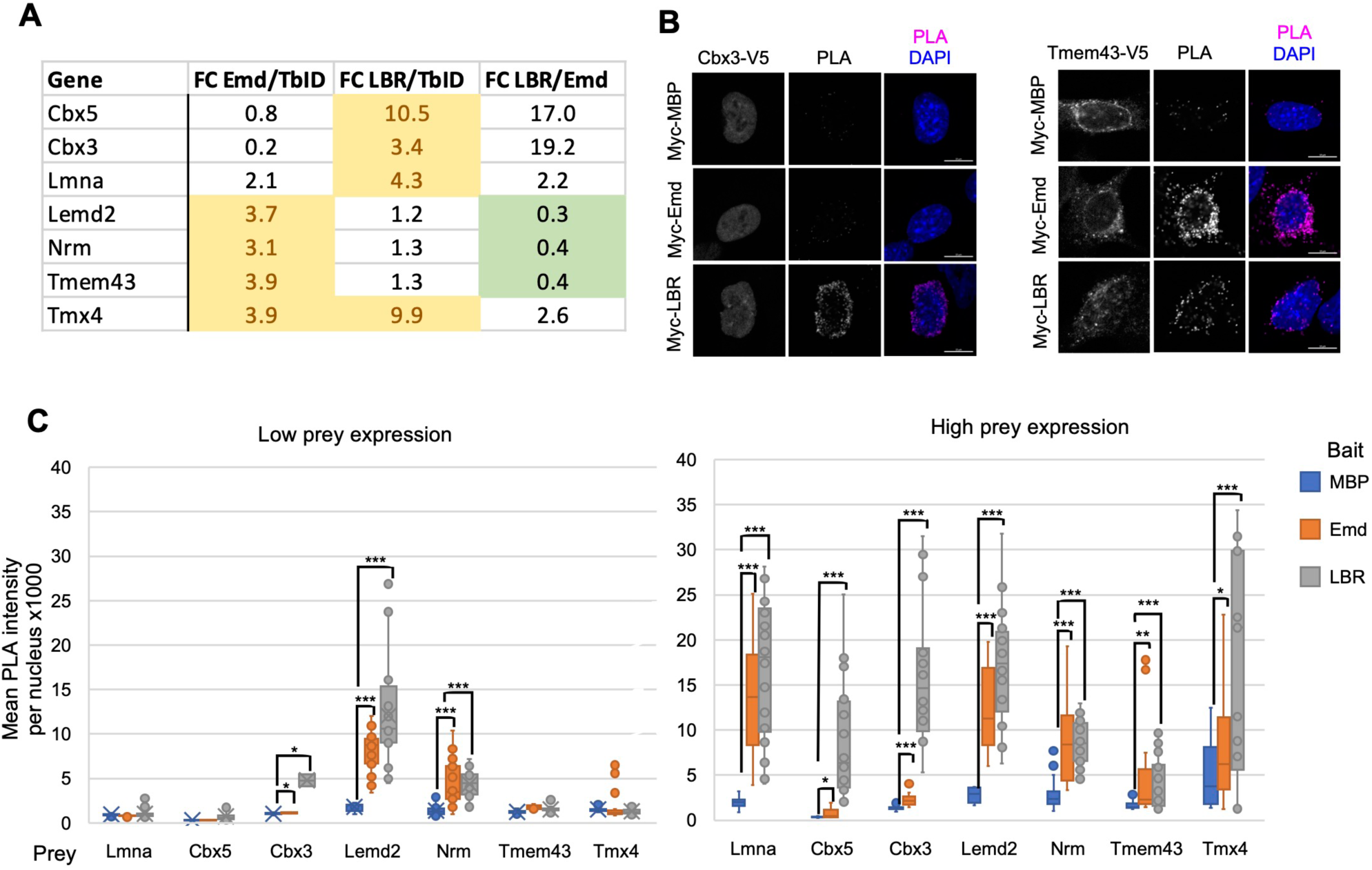
Proximity ligation analysis (PLA) of representative HCPP among well-characterized NE-associated proteins. (A) Summary of proximity labeling data by Emd-TbID and LBR-TbID, and results of the PLA analysis, from the group of HCPP analyzed. (B) Representative immunofluorescence images describing PLA obtained for cells stably expressing Myc-MBP, Myc-Emd, or Myc-LBR and transiently transduced with V5-tagged Cbx3 (left block of images) or Tmem43 (right block of images). First columns, anti-V5 staining; middle columns, PLA signal; right columns, merged imaged. Bars, 5 *µ*m. (C) Graphs depicting specific PLA signal obtained for samples in (A) with either low V5 expression (lower 50^th^ percentile) or high V5 expression (upper 50^th^ percentile). * p < 0.05, ** p < 0.01, *** p < 0.001.

**Figure 5.**
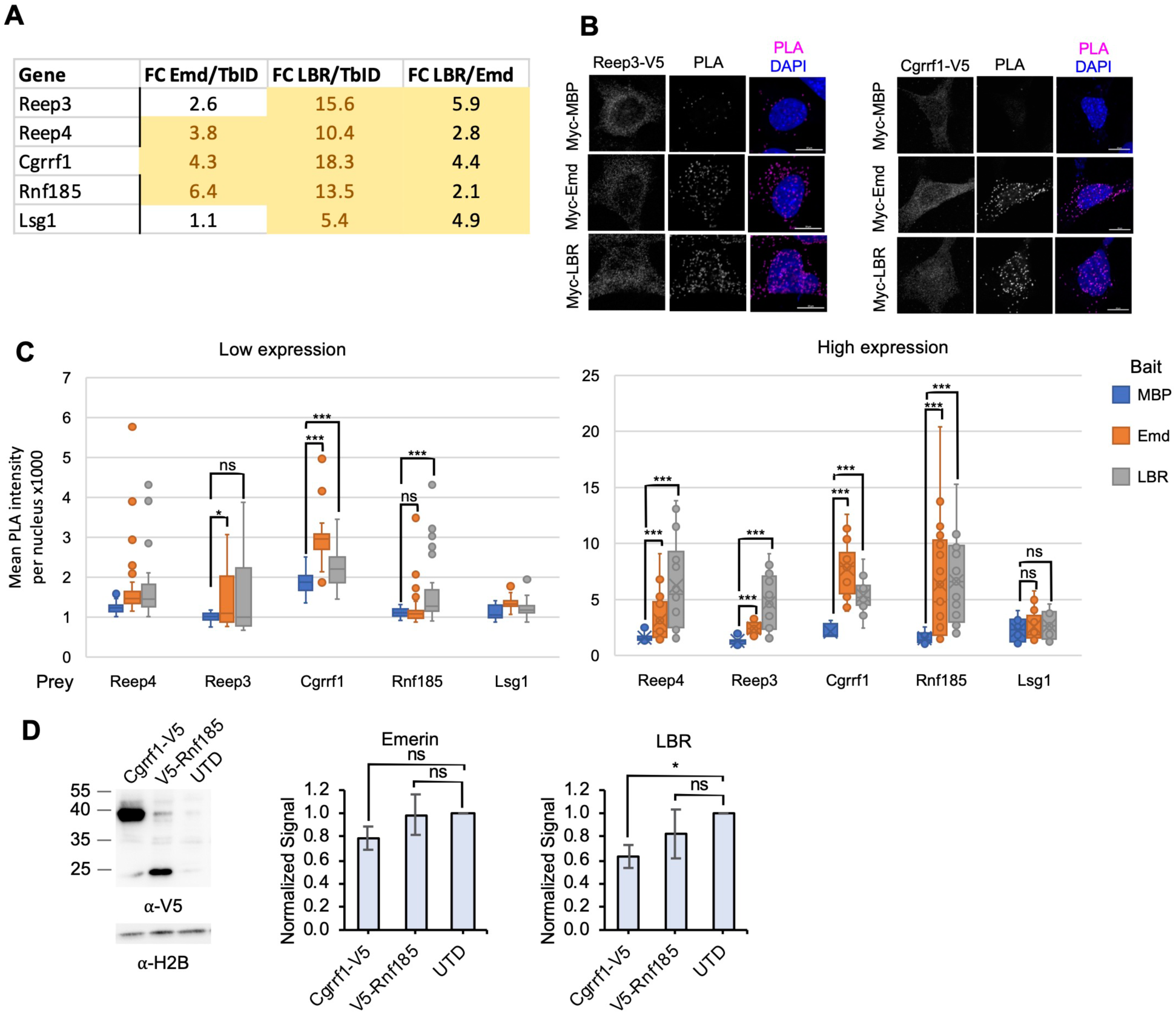
PLA of HCPP without known links to emerin or LBR. (A) Summary of proximity labeling data by Emd-TbID and LBR-TbID, and results of the PLA analysis, from HCPP analyzed. (B) Representative immunofluorescence images describing PLA obtained for cells stably expressing Myc-MBP, Myc-Emd, or Myc-LBR and transiently transduced with V5-tagged Reep3 (left block of images) or Cgrrf1 (right block of images). First columns, anti-V5 staining; middle columns, PLA signal; right columns, merged imaged. Bars, 5 *µ*m. (C) Graphs depicting specific PLA signal obtained for samples in (A) with either low V5 expression (lower 50^th^ percentile) or high V5 expression (upper 50^th^ percentile). * p < 0.05, ** p < 0.01, *** p < 0.001. (D) Left panel, Western blots documenting expression of V5-tagged Rnf185 and Cgrrf1 in stably transduced MEFs; right panels, quantification of the levels of endogenous emerin and LBR in these MEF strains, relative to untransduced cells (UTD), with normalization to α-tubulin. Original images used for quantification are in Fig. S6.

We first examined a group of prey with well-established NE localizations (Figs. 3 and 4). These included the LBR-binding heterochromatin components HP1α/Cbx5 and HP1γ*/*Cbx3, three emerin-linked INM proteins-lamin A, Lemd2 and Tmem43-and two proteins without known connections to either bait, Nrm and Tmx4. With low expression, only two prey-Lemd2 and Nrm-gave highly significant PLA signal with the LBR and Emd baits as compared to MBP (Fig. 4C). In the samples where PLA signal was detected only with high prey expression, both the LBR and Emd baits yielded significantly higher signal than MBP in all cases, regardless of the level of enrichment by LBR-TbID vs Emd-TbID (Fig. 4A). In some situations (i.e., Cbx3, Cbx5, Tmx4), the relative intensity of PLA signal with the Emd vs LBR baits was correlated with the level of proximity enrichment. Notably, the PLA signal for Cbx3/Cbx5 was much higher with the LBR bait than the emerin bait. In other cases, it was not: similar, significant PLA signal was obtained with both LBR-TbID and Emd-TbID for Lemd2, Nrm and Tmem43, even though these HCPP were enriched only with Emd-TbID.

Discrepancies between the results of proximity labeling and the PLA method are likely to reflect technical limitations associated with the PLA method. Since INM proteins evidently accumulate at the NE by binding to multiple partners with different affinities (see Introduction), prey analyzed by ectopic overexpression with the PLA method may populate a greater fraction of low affinity binding sites than prey expressed at endogenous levels, as detected by biotin proximity labeling. This could significantly affect the abundance of the different macromolecular interactions of prey detected by PLA. It also could bias epitope presentation for PLA detection, which could vary in different bait/prey macromolecular states. Furthermore, the PLA method is thought to have lower resolution (up to 30-40 nm) than biotin proximity labeling (10-20 nm) (May et al., 2020), thereby broadening the envelope for detectable proximity signal.

We subsequently analyzed five HCPP with unknown functional connections to emerin or LBR, that were not detectably concentrated at the NE (Figs. 3 and 5; Supporting Information Fig. S5). Four of these are TM proteins characterized as ER residents in UniProtKB. The fifth (Lsg1) is a non-membrane protein with large intranuclear and cytoplasmic pools that is implicated in nuclear export and cytoplasmic maturation of the large ribosomal subunit (Lo et al., 2010). The TM proteins selected for this analysis included Cgrrf1 (Glaeser et al., 2018; Lee et al., 2019) and Rnf185 (El Khouri et al., 2013; Glaeser et al., 2018; van de Weijer et al., 2020), ubiquitin E3 ligases implicated in ER-associated degradation (ERAD). These were enriched very highly with LBR-TbID (∼13-18-fold), and to a lesser degree by Emd-TbID (∼4-6-fold). As such, they could be involved in proteosomal degradation at the INM as described for certain E3 ligases in yeast (Natarajan et al., 2020).

In the low prey expression samples, a significant PLA signal was obtained for both LBR and emerin (with Cgrrf1) and for LBR with (Rnf185). At high prey expression, a significant PLA signal was obtained with both LBR and emerin for all prey except Lsg1 (i.e., Reep3, Reep4, Cgrrf1, and Rnf185). The accessibility of Lsg1 to proximity labeling by LBR and emerin but not to PLA detection could have multiple explanations (2^nd^ paragraph, above). In summary, the PLA results obtained with these relatively uncharacterized prey, together with the results for previously characterized NE proteins, validate the neighborhood assignments obtained by TbID labeling in 11 out of 12 instances. This suggests that the great majority of proteins in our HCPP list have physical proximity to emerin and/or LBR at the NE.

In an initial functional analysis, we examined Cgrrf1 and Rnf185 for a potential role in the proteosomal turnover of emerin and LBR, proteins that have half-lives of ∼1.5-3.5 days in myoblasts (Buchwalter et al., 2019). First, we analyzed the levels of endogenous emerin and LBR in MEFs that were stably transduced with the ectopic E3 ligases (Fig. 5D). In cells overexpressing Cgrrf1 (Fig. 5D, left panel), we observed a significant decrease in the level of LBR but no detectable change in the level of emerin (Fig. 5D, right panels). Conversely, no changes in either LBR or emerin were detected in cells overexpressing Rnf185 (Fig. 5D). In a complementary approach, we analyzed MEFs in which Cgrrrf1 or Rnf185 were depleted by RNAi to ∼80% at the mRNA level (Supporting Information Fig. S7). In these cases, no differences in the levels of endogenous LBR or emerin were detected as compared to control RNAi. The reduction in LBR levels seen with overexpression of Cgrrf1 suggests a potential role for this E3 ligase in turnover of LBR. Conversely, the lack of a detectable effect with Cgrrf1 knockdown could be due to incomplete depletion of the latter, or could reflect the existence of additional compensatory E3 ligases that help to maintain steady state levels of LBR with reduced Cgrrf1. Regardless, the very strong enrichment of Cgrrf1 and Rnf185 with LBR-TbID and Emd-TbID, as compared to the much lower labeling of other ER-localized E3 ligases detected in our datasets (Supporting Information Table S2), argue that further analysis of these E3 ligases is warranted to query their potential role in regulating NE functions and/or protein levels.

## Discussion

Here we investigated the neighborhoods of emerin and LBR in MEFs using TbID-based proximity labeling and quantitative proteomics. By comparing prey enrichment patterns obtained with emerin and LBR baits to those of unfused TbID, we generated a HCPP list that included a cohort of well characterized NE components and many additional proteins that heretofore have not been functionally connected to emerin or LBR. We used the PLA as an orthogonal approach to query proximity relationships of emerin and LBR at the NE to selected HCPP, including both NE-concentrated proteins and components with no apparent NE enrichment. These experiments confirmed NE proximity for 11 of the 12 HCPP prey tested, supporting the spatial relationships suggested by our TbID datasets. Overall, our proximity labeling approach revealed both shared and distinctive HCPP for LBR and emerin.

How can the labeling of NE-concentrated prey be interpreted in the context of NE organization? In the simplest model, preferential enrichment with either the LBR or emerin baits may reflect the presence of certain prey in compositionally distinct macromolecular complexes containing either LBR or emerin. Consistently, some of the NE proteins that were enriched with either LBR or emerin with strong preference have been selectively linked to the corresponding protein by biochemical and cell-based studies (see Results). Conversely, NE-associated HCPP that were strongly enriched with both the LBR and emerin baits may reflect the presence of these prey in compositionally overlapping macromolecular complexes containing LBR or emerin. However, in some cases strong prey labeling by both LBR and emerin may arise by default due to the concentration of LBR, emerin and prey in a spatially constrained sub-domain(s) of the INM. One such INM subdomain is the membrane juxtaposed to lamin filaments, which comprises only a small portion of the total INM surface (de Leeuw et al., 2018). Since emerin, LBR and many other INM proteins directly interact with lamins (Pawar and Kutay, 2021), dynamic binding and dissociation of these proteins from lamin filaments could stochastically position bait/prey pairs within an effective distance for proximity labeling even when they are not functionally complexed. The presence of long intrinsically disordered regions in the nucleoplasmic domains of LBR, emerin and other NE proteins could further diminish the resolution of proximity labeling. These intrinsic limitations merit consideration in future studies.

A major fraction of the HCPP of emerin and LBR are localized throughout the peripheral ER. Enrichment of certain of these prey with our probes may reflect functions of emerin or LBR at ER regions other than the NE. Alternatively, ER-localized HCPP may have functionally significant associations with emerin or LBR at the NE, since most peripheral ER proteins have access to both the ONM and INM and can rapidly flux between these membranes (Ungricht and Kutay, 2015). Supporting this possibility, the HCPP included a number of ER-localized TM proteins that have been linked to NE functions or dynamics including Ankle2 (Asencio et al., 2012), Reep3/4 (Schlaitz et al., 2013), Lunapark (Casey et al., 2015; Hirano et al., 2020) and Dnajb12 (Goodwin et al., 2014). Moreover, our PLA detected Reep3 and Reep4 in the proximity of LBR and emerin at the NE. In this regard, the ER contains at least 21 membrane-embedded ubiquitin E3 ligases (Fenech et al., 2020), but only three of these, Cgrrf1 and Rnf185 (and to a lesser extent Bfar), were strongly enriched by LBR and emerin. Considering our initial evidence that Cgrrf1 can modulate the level of endogenous LBR, these E3 ligases merit further investigation in the context of INM homeostasis.

In contrast to the HCPP identified by emerin or LBR that are localized to the ER network, a substantial fraction have been localized to mitochondria or to distal membranes in the secretory pathway such as Golgi. In these cases, the prey may be labeled by a peripheral ER pool of emerin-TbID or LBR-TbID that might be present at ER-organelle membrane contact sites (MCS). In addition, since emerin is known to traverse the secretory pathway *en route* to the plasma membrane and endosome (Buchwalter et al., 2019), labeling could occur in secretory pathway compartments downstream of the peripheral ER. Potential functions of these prey in relation to emerin and LBR, either at the NE, MCS, or other cellular locations, remain to be investigated.

Proximity labeling approaches have been used by others to investigate interactions of emerin in cultured cells (Go et al., 2021; Moser et al., 2020; Muller et al., 2020), albeit with different enzymes (BioID and APEX2) and cell types (HEK293, HeLa and U2OS) from those used in our analysis. Our HCPP list contains a minor fraction of the statistically significant emerin prey identified in this other work (Supporting Information, Table S3). These included 43 of the 290 interacting proteins reported by Go et al (Go et al., 2021), 9 of the 56 prey reported by Müller et al (Muller et al., 2020) and 3 of the 44 high confidence interactors reported by Moser et al (Moser et al., 2020). The disparities between these studies may be explained by differences in the experimental systems, cultured cell types and analytical methods.

The two emerin prey that were identified by all four proximity studies-TOR1AIP1 and VAPA-are noteworthy. TOR1AIP1 is involved in diverse aspects of NE functions (Rampello et al., 2020), and has been physically and functionally linked to emerin (Shin et al., 2013). VAPA is a member of the VAP family of ER proteins involved in the formation of MCS between the ER and other organelles via a “FFAT” peptide motif in interacting proteins (James and Kehlenbach, 2021). Intriguingly, emerin contains a conserved region (aa 92-97 in human emerin, DDYYEE) that closely resembles FFAT motif variants (Di Mattia et al., 2020). This suggests a potential function for emerin at MCS by interaction with VAPA or with other VAP family members such as VAPB, which partially resides in the INM (James et al., 2019), or Mospd1, identified as an emerin HCPP in our analysis. However, whether VAPA is linked functionally to emerin and/or to TOR1AIP1 remains to be determined.

In summary, our analysis has identified new potential functional partners of emerin and LBR, and additionally, identifies proteins that may be concentrated at the INM or that rapidly flux through this compartment. Our comparative analysis of emerin and LBR using quantitative proteomics highlights the distinctive properties of these INM proteins. In future work, application of similar quantitative methods to a large cohort of INM proteins should permit a broad-ranging analysis of local environments of the INM. Critical assessment of labeling patterns may be further enhanced by adjusting the expression level of proximity labeling probes, and/or by fusing biotinylating enzymes to INM proteins at the genomic level to circumvent complications due to ectopic expression. In combination with additional tools, such as light microscopy-based proximity analysis such as FRET/FLIM, this could lay the foundation for a comprehensive evaluation of the landscape of the INM and how this changes in different cellular states.

## Data Availability

Raw proteomic datasets are available in MassIVE and ProteomeXchange with the following accession numbers: MSV000089356 and PXD033571.

## Author Contributions

LG directed the project and wrote the manuscript. JRY provided equipment and support personnel for MS analysis. L-CC, XZ and KA designed subsets of the experiments. XZ prepared and analyzed proteomics samples. KA prepared TbID cell strains and samples. L-CC and JAN implemented the proximity ligation analysis. JAN carried out the RNAi. SM-B and L-CC did the statistical analysis. SB did bioinformatic analysis and helped with data interpretation and figure preparation. TCB and AYT provided the TbID clone. EL, JAN and KA implemented the molecular cloning.

## Notes

The authors declare no competing financial interests.

## Supporting Information

**Figure S1.**
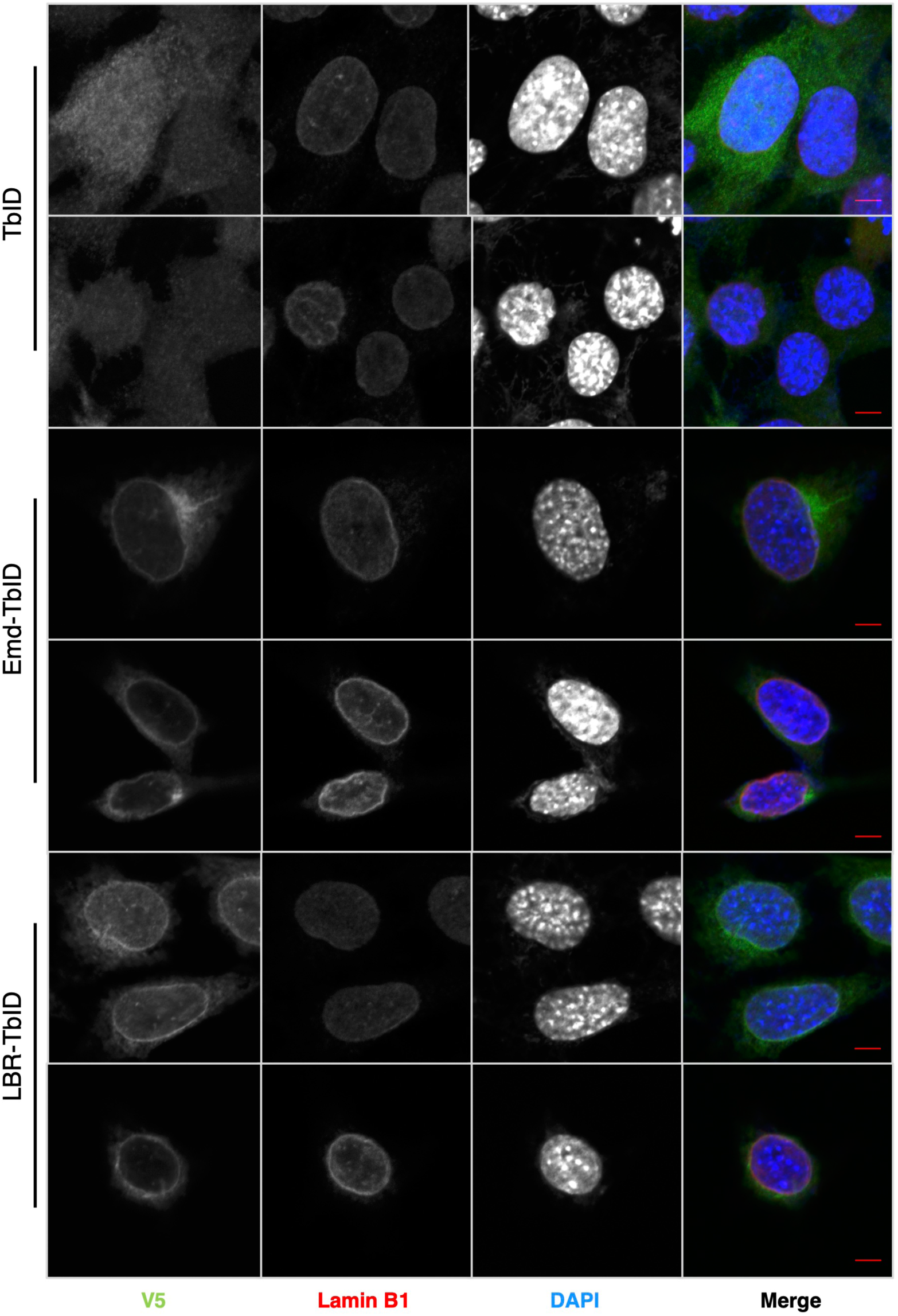
Gallery of light micrographs showing localization of TbID probes in MEFs. Stably transduced MEFs were fixed and labeled for immunofluorescence as in Fig. 1, to detect V5-tagged TbID constructs, endogenous lamin B1 and DNA as indicated. Bars, 5 *µ*m.

**Figure S2.**
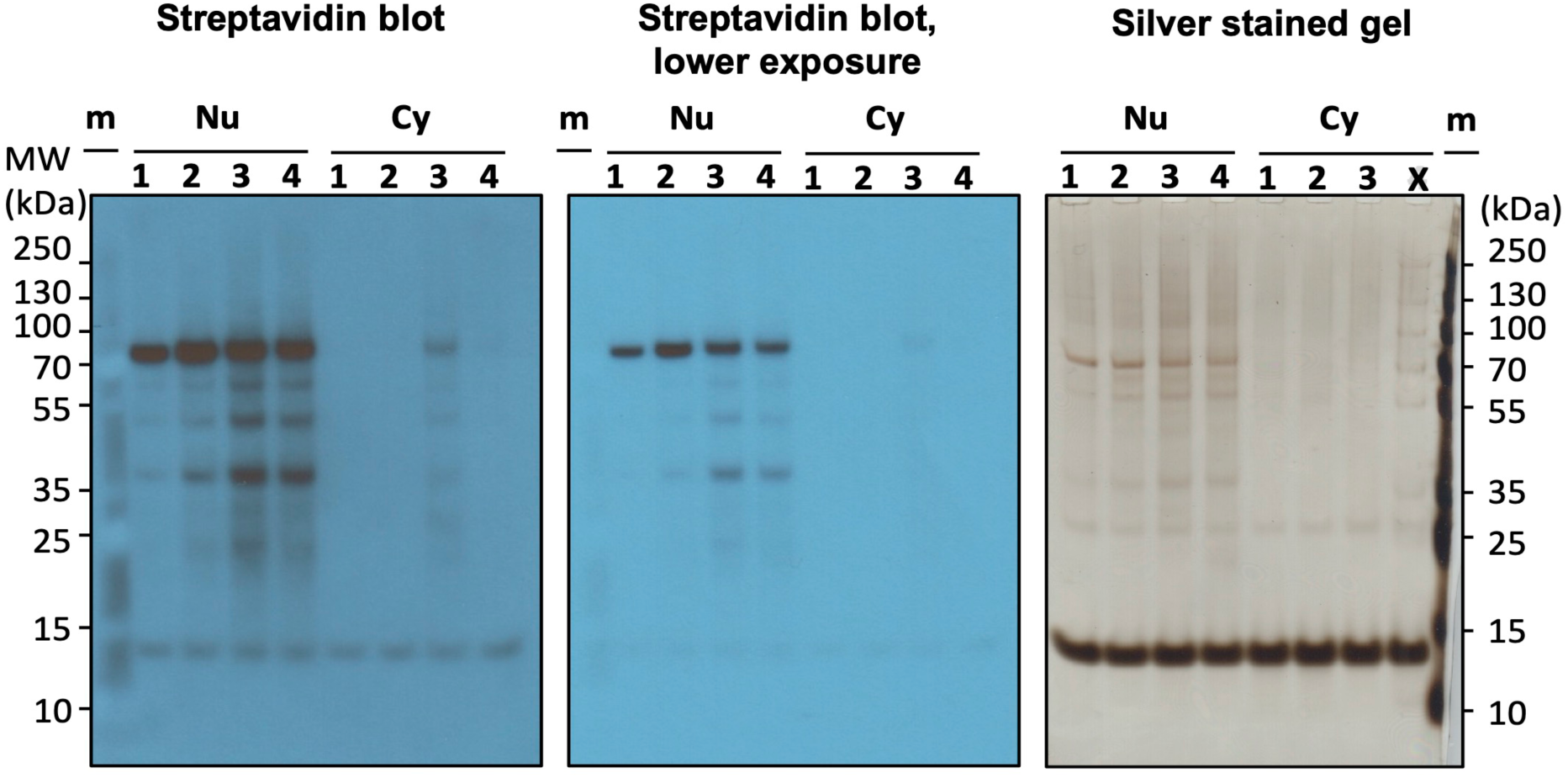
Time course of biotin labeling in MEFs expressing emerin-TbID. Cells were incubated with 500 *µ*M biotin (lanes 1, 2, 4) or 50 *µ*M biotin (lanes 3) for 10 min (lanes 1), 1 h (lanes 2) or 2 h (lanes 3, 4), and cells were homogenized and fractionated to yield a low speed nuclear pellet (Nu) and a postnuclear supernatant (Cy). After sample solubilization in SDS, proteins were captured on streptavidin beads. A portion of the samples was eluted from beads by boiling in SDS/urea, and analyzed by SDS-PAGE. Left two panels, Western blots of gels probed with HRP-streptavidin. Right, silver stained gel to validate equivalent sample loading. m lanes, molecular weight markers.

**Figure S3.**
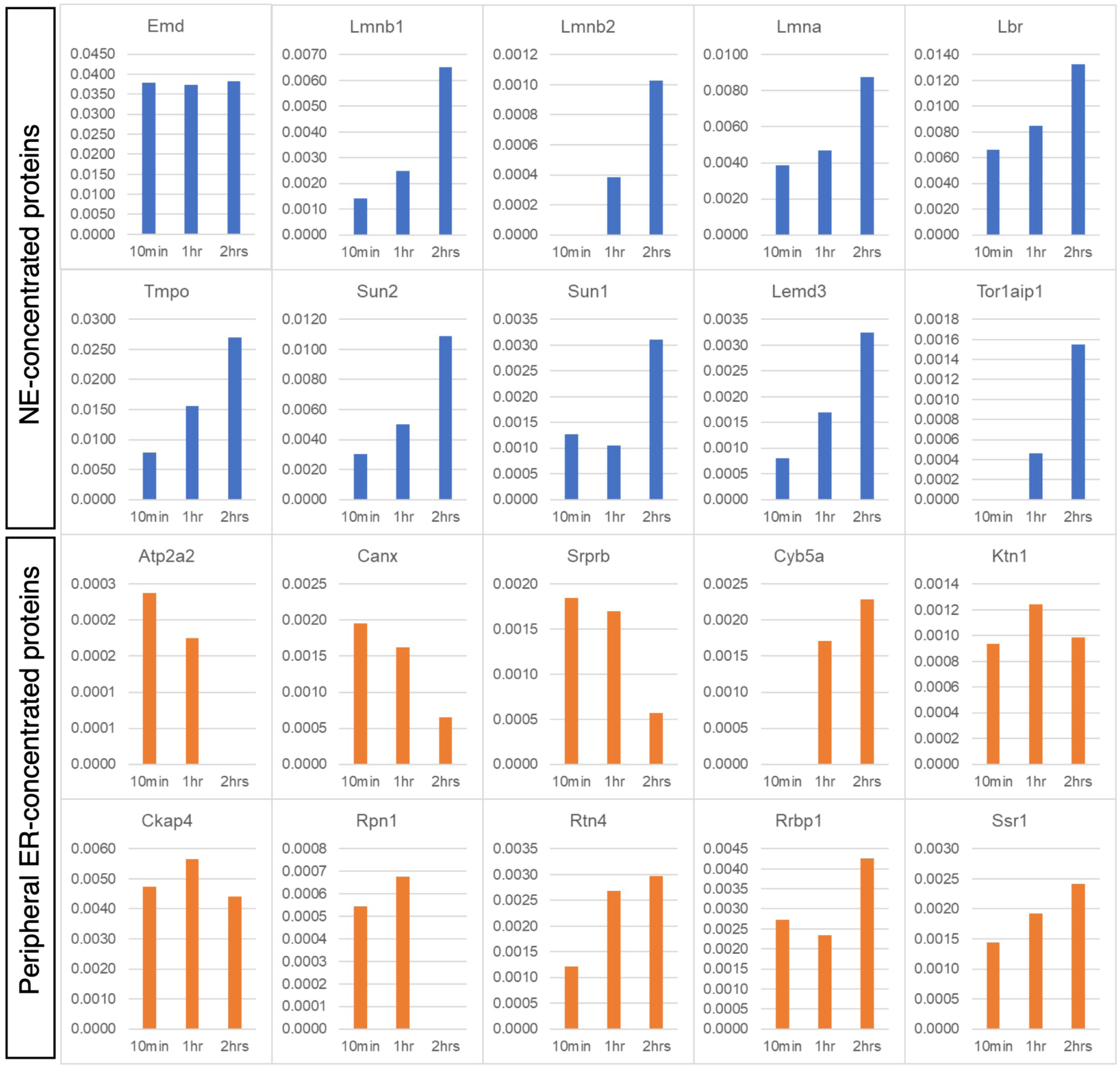
Time course of biotin labeling in MEFs expressing emerin-TbID. Cells were incubated with biotin for 10 min, 1 h or 2 h, as indicated, and biotinylated proteins were captured on streptavidin beads under denaturing conditions. Subsequently, samples were processed by trypsin digestion and dimethyl labeling and analyzed by LC-MS-MS, allowing comparison of 3 samples in a single MS run (Experimental Procedures and Table S1). Graphs show NSAF values for benchmark proteins of the NE and peripheral ER captured at the 3 time points.

**Figure S4.**
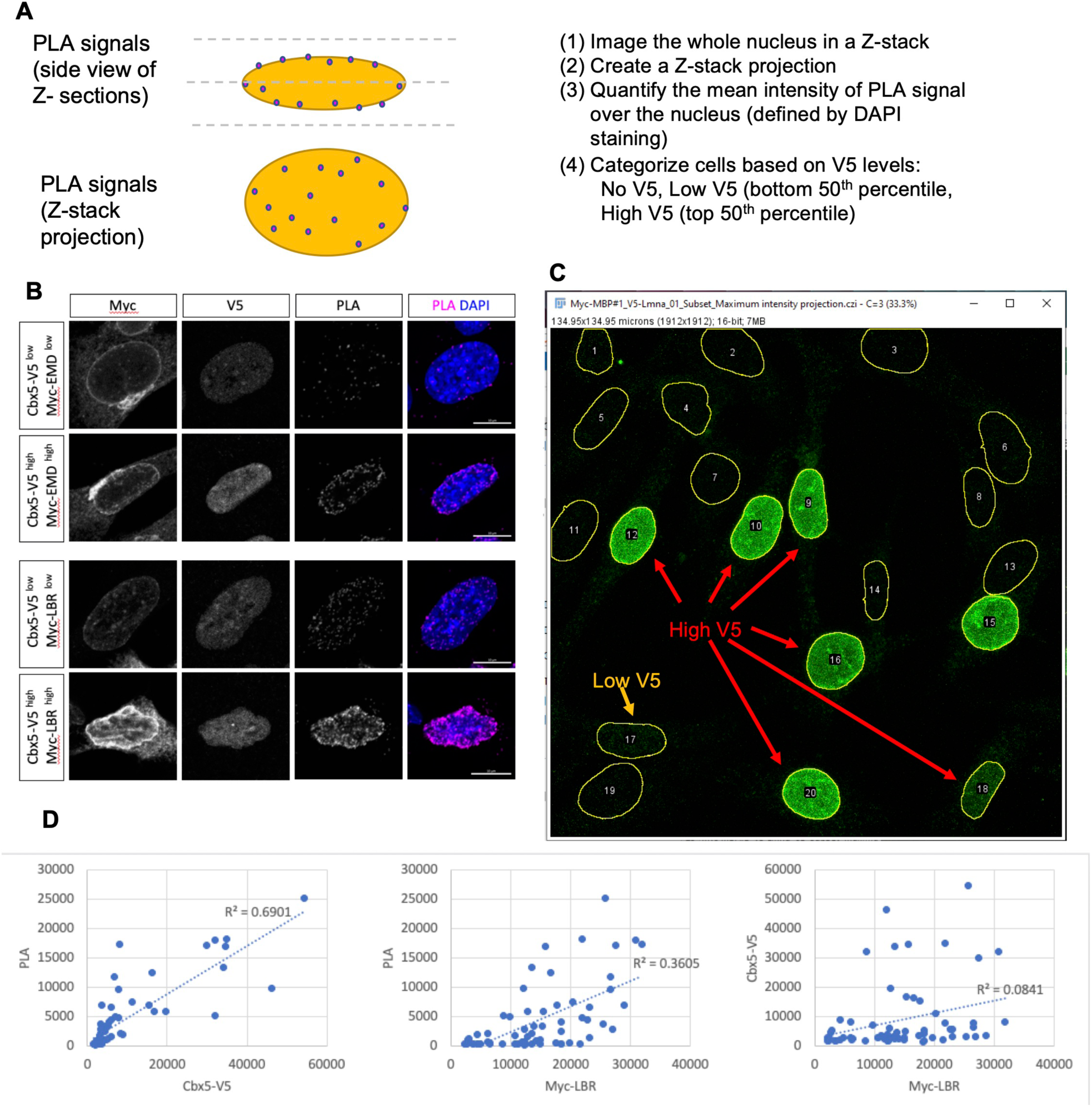
Quantification of PLA signal at the NE/nucleus with bait-prey pairs. (A) Strategy for quantification. Left panel: Cartoon illustrating results of a PLA reaction in a cell. Individual PLA foci are depicted by dots. Due to the flatness of the nucleus in MEFs, it was not possible to consistently obtain Z-sections that imaged only the nuclear rim without containing dorsal and ventral tangential areas of the NE. Right panel: Steps in analysis. Sequential Z-sections containing the DAPI-stained nucleus were collected and projected as a Z-stack. The mean nuclear PLA intensity in the Z-stack was then calculated for a group of ∼10-30 cells, together with the intensity of V5-tagged bait (and in some cases, the intensity of Myc-tagged prey). Based on V5-prey expression, cells were sorted as “low expression” (lower 50^th^ percentile) or “high expression (upper 50^th^ percentile) groups and separately graphed. (B) Immunofluorescence micrographs of cells expressing V5-tagged Cbx5 and Myc-tagged LBR or emerin and analyzed by PLA, shown for low and high expression of Cbx5 and Myc. (C) Field of cells showing examples of PLA results in cells with low or high V5-Cbx5 and Myc-LBR expression. (D) Graphs depicting nuclear PLA intensity in individual cells as a function of the level of V5-Cbx5 and Myc-LBR. The PLA intensity was strongly correlated with the level of V5-Cbx-5 (prey) expression (R^2^, 0.69) and moderately correlated with the Myc-LBR (bait) expression (R^2^, 0.36). However the level of V5-prey and Myc-bait were uncorrelated (R^2^, 0.08).

**Figure S5.**
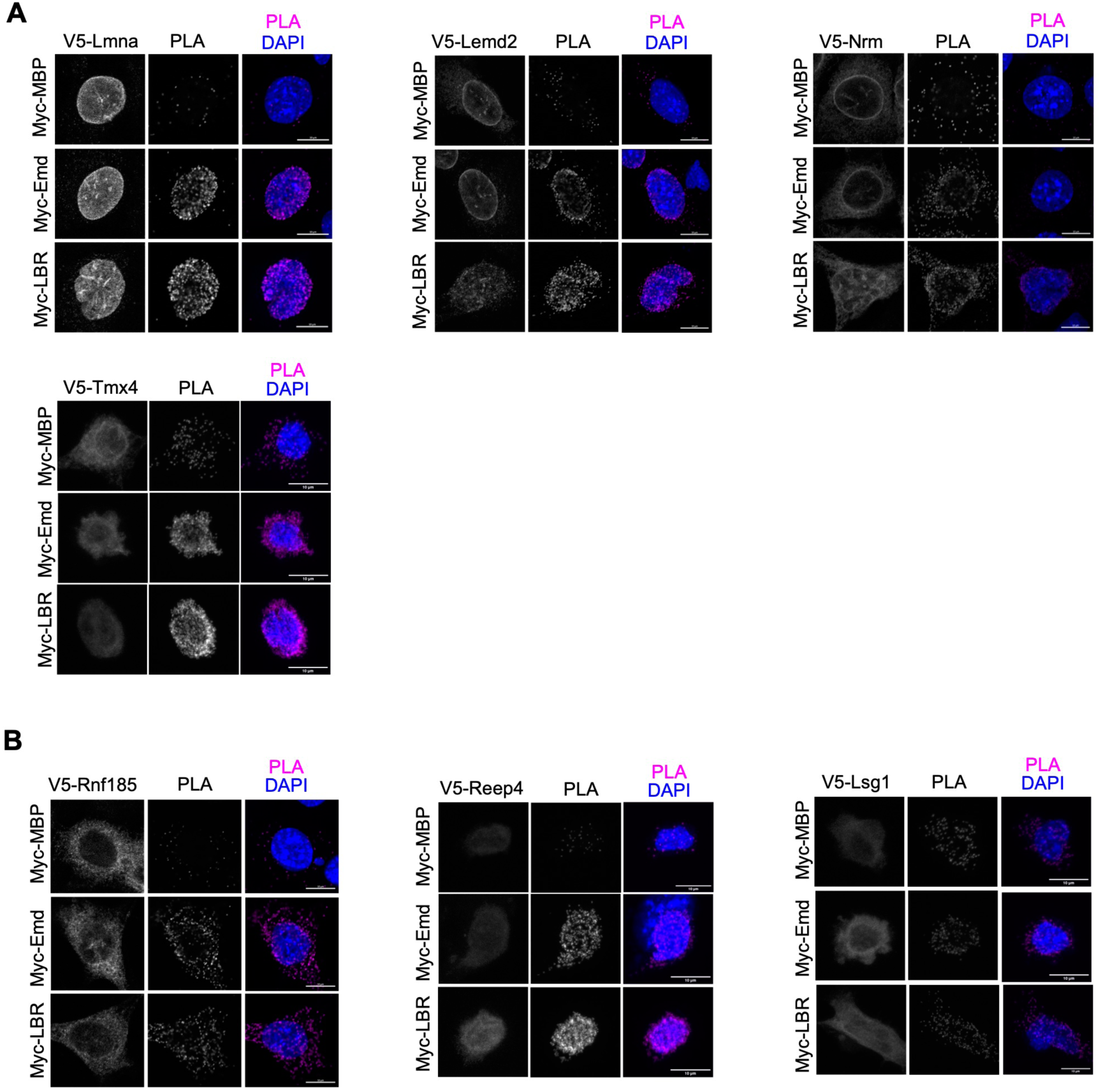
Galleries of light micrographs of typical cells used to quantify nuclear PLA with different bait-prey pairs. (A) Examples of cells expressing benchmark NE prey that are not shown in Fig 4B. Results from cell populations involving these prey are graphed in Fig. 4C. (B) Examples of cells expressing exemplary HCPP not previously linked to emerin or LBR and not shown in Fig. 5B. Results from cell populations with these prey are graphed in Fig 5C.

**Figure S6.**
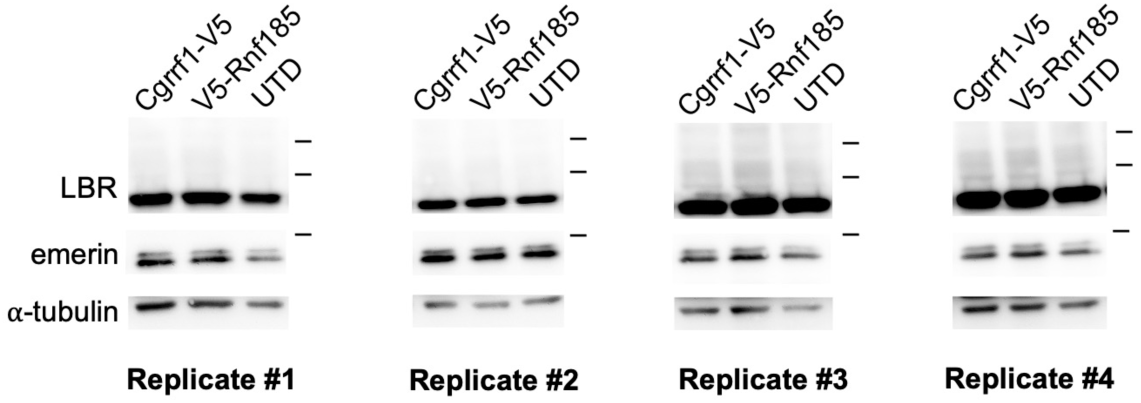
Images of Western blots of the replicates used to calculate the level of endogenous emerin and LBR in MEFs overexpressing Cgrrf1 or Rnf185. Intensities for emerin/LBR were normalized to α-tubulin. Data is plotted in Fig. 5D.

**Figure S7.**
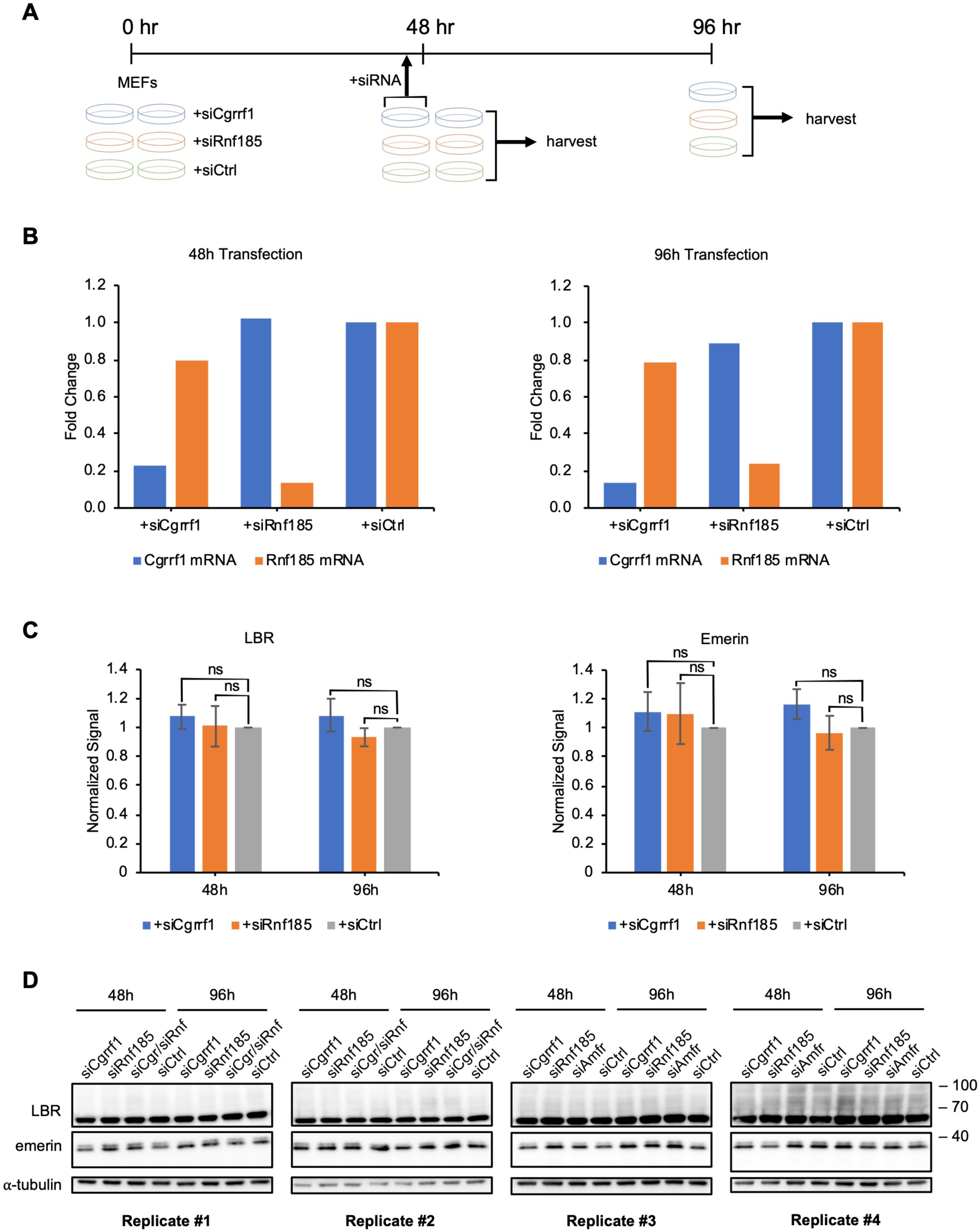
Analysis of endogenous emerin and LBR in cells with knockdown of Rnf185 or Cgrrf1. (A) Scheme for RNAi analysis. Cells were transfected with siRNA pools to target Rnf185 or Cgrrf1, or with a control siRNA pool. 48 h later, one set of samples was harvested for Western blot analysis, and a second set was transfected with another round of siRNA. Cells from the second set were harvested for Western analysis after further 48 hr. (B) Levels of mRNA for Rnf185 and Cgrrf1 in the various RNAi samples (Fig. S6A), as determined by q-RT-PCR and normalization to the levels in control siRNA cells. In all cases, efficient mRNA depletion was obtained. (C) Quantification of levels of emerin and LBR in cells with the various RNAi conditions. (D) Images of the Western blots (4 replicates) used to quantify emerin/LBR levels in Fig. S6C.

